## Supplemental Table 2 for "A stable, distributed code for cue value in mouse cortex during reward learning"

| **Cue decoding performance, Bonferroni corrected p-values for group comparisons (Fig. 3G)** | | | | | | | | | | | | |
| --- | --- | --- | --- | --- | --- | --- | --- | --- | --- | --- | --- | --- |
| **Time from cue ->** | -0.25 | 0 | 0.25 | 0.5 | 0.75 | 1 | 1.25 | 1.5 | 1.75 | 2 | 2.25 | 2.5 |
| Value>baseline |  | 1 | 1 | 1 | 2.61E-19 | 5.39E-71 | 3.97E-99 | 1.08E-90 | 5.01E-96 | 3.70E-94 | 4.85E-101 | 1.00E-97 |
| Value like > baseline |  | 1 | 1 | 1 | 3.96E-25 | 1.86E-87 | 1.14E-102 | 8.10E-107 | 1.61E-99 | 1.33E-107 | 2.22E-109 | 7.68E-103 |
| Untuned > baseline |  | 1 | 1 | 1 | 2.61E-19 | 5.39E-71 | 3.97E-99 | 1.08E-90 | 5.01E-96 | 3.70E-94 | 4.85E-101 | 1.00E-97 |
| Value>value like | 1 | 1 | 1 | 1 | 1 | 2.70E-05 | 2.53E-13 | 1.52E-08 | 8.42E-13 | 8.92E-10 | 2.75E-12 | 1.02E-11 |
| Value>untuned | 1 | 1 | 1 | 1 | 1.43E-06 | 1.13E-34 | 2.55E-44 | 1.34E-39 | 1.16E-46 | 1.62E-48 | 3.15E-43 | 2.00E-29 |
| Value like>untuned | 1 | 1 | 1 | 1 | 0.00107 | 6.16E-20 | 4.00E-17 | 3.00E-19 | 1.53E-19 | 3.94E-25 | 1.89E-17 | 5.53E-08 |
| **Cue decoding (population), Bonferroni corrected p-values for group comparisons (Fig. 3H)** | | | | | | | | | | | | |
| **Pseudoensemble ->** | 1 | 5 | 10 | 25 | 50 | 75 | 100 | 200 |  |  |  |  |
| Value>chance | 0.012 | 0.001 | 0.001 | 0.001 | 0.001 | 0.001 | 0.001 | 0.001 |  |  |  |  |
| Value like>chance | 0.019 | 0.001 | 0.001 | 0.001 | 0.001 | 0.001 | 0.001 | 0.001 |  |  |  |  |
| Untuned>chance | 0.102 | 0.001 | 0.001 | 0.001 | 0.001 | 0.001 | 0.001 | 0.001 |  |  |  |  |
| Value>value like | 1 | 1 | 1 | 1 | 1 | 1 | 1 | 1 |  |  |  |  |
| Value>untuned | 1 | 0.44 | 0.32 | 0.1428 | 0.123 | 0.1662 | 0.2028 | 0.726 |  |  |  |  |
| Value like>untuned | 1 | 0.86 | 0.62 | 0.1896 | 0.05712 | 0.02832 | 0.021 | 0.05928 |  |  |  |  |
| Value<value like | 1 | 1 | 1 | 1 | 1 | 1 | 0.918 | 0.828 |  |  |  |  |
| Value<untuned | 1 | 1 | 1 | 1 | 1 | 1 | 1 | 1 |  |  |  |  |
| Value like<untuned | 1 | 1 | 1 | 1 | 1 | 1 | 1 | 1 |  |  |  |  |
| **Value decoding (population), Bonferroni corrected p-values for group comparisons (Fig. 3I)** | | | | | | | | | | | | |
| **Pseudoensemble ->** | 1 | 5 | 10 | 25 | 50 | 75 | 100 | 200 |  |  |  |  |
| Value>chance | 0.03 | 0.001 | 0.001 | 0.001 | 0.001 | 0.001 | 0.001 | 0.001 |  |  |  |  |
| Value like>chance | 0.031 | 0.001 | 0.001 | 0.001 | 0.001 | 0.001 | 0.001 | 0.001 |  |  |  |  |
| Untuned>chance | 0.126 | 0.001 | 0.001 | 0.001 | 0.001 | 0.001 | 0.001 | 0.001 |  |  |  |  |
| Value>value like | 1 | 0.72 | 0.27 | 0.052 | 0.0162 | 0.0108 | 0.00642 | 7.02E-04 |  |  |  |  |
| Value>untuned | 1 | 0.28 | 0.15 | 0.0068 | 6.00E-06 | 1.20E-05 | 6.00E-06 | 6.00E-06 |  |  |  |  |
| Value like>untuned | 1 | 1 | 1 | 1 | 1 | 1 | 0.93 | 1 |  |  |  |  |
| Value<value like | 1 | 1 | 1 | 1 | 1 | 1 | 1 | 1 |  |  |  |  |
| Value<untuned | 1 | 1 | 1 | 1 | 1 | 1 | 1 | 1 |  |  |  |  |
| Value like<untuned | 1 | 1 | 1 | 1 | 1 | 1 | 1 | 1 |  |  |  |  |

**Supplemental Table 2**. Bootstrapped distributions (1000 samples), *p*-values from directional overlap (related to Fig. 3).
