## Supplemental Table 3 for "A stable, distributed code for cue value in mouse cortex during reward learning"

| **Value cells, Bonferroni corrected p-values for pairwise region comparisons (Fig. 4A, S9A-B left)** | | | | | | | | | |
| --- | --- | --- | --- | --- | --- | --- | --- | --- | --- |
|  | ALM | ACA | FRP | PL | ILA | ORB | DP | TTd | AON |
| ALM |  | 0.0365 | 1 | 0.22 | 1 | 0.0282 | 1 | 1 | 1 |
| ACA | 0.0365 |  | 1 | 1 | 1 | 1 | 0.64 | 0.14 | 0.30 |
| FRP | 1 | 1 |  | 1 | 1 | 1 | 1 | 1 | 1 |
| PL | 0.22 | 1 | 1 |  | 1 | 1 | 1 | 0.90 | 1 |
| ILA | 1 | 1 | 1 | 1 |  | 1 | 1 | 1 | 1 |
| ORB | 0.0282 | 1 | 1 | 1 | 1 |  | 1 | 0.50 | 0.74 |
| DP | 1 | 0.64 | 1 | 1 | 1 | 1 |  | 1 | 1 |
| TTd | 1 | 0.14 | 1 | 0.90 | 1 | 0.50 | 1 |  | 1 |
| AON | 1 | 0.30 | 1 | 1 | 1 | 0.74 | 1 | 1 |  |
|  | Motor | PFC | Olf. |  |  |  |  |  |  |
| Motor |  | 0.0062 | 1 |  |  |  |  |  |  |
| PFC | 0.0062 |  | 0.0002 |  |  |  |  |  |  |
| Olfactory | 1 | 0.0002 |  |  |  |  |  |  |  |
| **Value-like cells, Bonferroni corrected p-values for pairwise region comparisons (Fig. 4B, S9A-B right)** | | | | | | | | | |
|  | ALM | ACA | FRP | PL | ILA | ORB | DP | TTd | AON |
| ALM |  | 1 | 1 | 0.26 | 0.36 | 1.10E-05 | 0.08 | 0.0002 | 0.0407 |
| ACA | 1 |  | 1 | 1 | 1 | 1 | 1 | 1 | 1 |
| FRP | 1 | 1 |  | 1 | 1 | 0.0443 | 1 | 0.0494 | 0.67 |
| PL | 0.26 | 1 | 1 |  | 1 | 0.36 | 1 | 0.22 | 1 |
| ILA | 0.36 | 1 | 1 | 1 |  | 1 | 1 | 0.89 | 1 |
| ORB | 1.10E-05 | 1 | 0.0443 | 0.36 | 1 |  | 1 | 1 | 1 |
| DP | 0.08 | 1 | 1 | 1 | 1 | 1 |  | 1 | 1 |
| TTd | 0.0002 | 1 | 0.0494 | 0.22 | 0.89 | 1 | 1 |  | 1 |
| AON | 0.04 | 1 | 0.67 | 1 | 1 | 1 | 1 | 1 |  |
|  | Motor | PFC | Olf. |  |  |  |  |  |  |
| Motor |  | 2.34E-05 | 1.17E-07 |  |  |  |  |  |  |
| PFC | 2.34E-05 |  | 0.0443 |  |  |  |  |  |  |
| Olfactory | 1.17E-07 | 0.0443 |  |  |  |  |  |  |  |
| **Population decoding of value, region comparisons (bootstrap, Bonferroni corrected p-values) (Fig. 4E)** | | | | | | | | | |
|  | ALM | ACA | FRP | PL | ILA | ORB | DP | TTd | AON |
| ALM |  | 1 | 1 | 1 | 1 | 1 | 1 | 1 | 1 |
| ACA | 1 |  | 1 | 1 | 1 | 1 | 1 | 1 | 1 |
| FRP | 1 | 1 |  | 1 | 1 | 1 | 1 | 1 | 1 |
| PL | 1 | 1 | 1 |  | 1 | 1 | 1 | 1 | 1 |
| ILA | 1 | 1 | 1 | 1 |  | 1 | 1 | 1 | 1 |
| ORB | 1 | 1 | 1 | 1 | 1 |  | 1 | 1 | 1 |
| DP | 1 | 1 | 1 | 1 | 1 | 1 |  | 1 | 1 |
| TTd | 1 | 1 | 1 | 1 | 1 | 1 | 1 |  | 1 |
| AON | 1 | 1 | 1 | 1 | 1 | 1 | 1 | 0.680148 |  |
|  | Motor | PFC | Olf. |  |  |  |  |  |  |
| Motor |  | 1 | 1 |  |  |  |  |  |  |
| PFC | 0.23 |  | 0.74 |  |  |  |  |  |  |
| Olfactory | 0.70 | 1 |  |  |  |  |  |  |  |

**Supplemental Table 3**. Generalized linear mixed-effects model, *p*-values from region contrasts (related to Fig. 4 and Fig. S9).
