## Supplemental Table 4 for "A stable, distributed code for cue value in mouse cortex during reward learning"

| **Slope, Bonferroni corrected p-values for pairwise group comparisons (Fig. 5H)** | | | | | | | | | |
| --- | --- | --- | --- | --- | --- | --- | --- | --- | --- |
|  | history | value | value-like | untuned | licks |  |  |  |  |
| history |  | 1.00E-08 | 1.00E-08 | 1.00E-08 | 0.0062 |  |  |  |  |
| value | 1 |  | 0.01 | 0.92 | 1 |  |  |  |  |
| value-like | 1 | 0.99 |  | 1.00 | 1 |  |  |  |  |
| untuned | 1 | 0.0803 | 0.0033 |  | 1.00 |  |  |  |  |
| licks | 0.99 | 1.00E-08 | 1.00E-08 | 0.0003 |  |  |  |  |  |
| **History cells, Bonferroni corrected p-values for pairwise region comparisons (Fig. 5I)** | | | | | | | | | |
|  | ALM | ACA | FRP | PL | ILA | ORB | DP | TTd | AON |
| ALM |  | 1 | 1 | 1 | 1 | 1 | 1 | 1 | 1 |
| ACA | 1 |  | 1 | 1 | 1 | 1 | 0.66 | 1 | 1 |
| FRP | 1 | 1 |  | 1 | 1 | 1 | 1 | 1 | 1 |
| PL | 1 | 1 | 1 |  | 1 | 1 | 1 | 1 | 1 |
| ILA | 1 | 1 | 1 | 1 |  | 1 | 1 | 1 | 1 |
| ORB | 1 | 1 | 1 | 1 | 1 |  | 1 | 1 | 1 |
| DP | 1 | 0.66 | 1 | 1 | 1 | 1 |  | 1 | 1 |
| TTd | 1 | 1 | 1 | 1 | 1 | 1 | 1 |  | 1 |
| AON | 1 | 1 | 1 | 1 | 1 | 1 | 1 | 1 |  |
|  | Motor | PFC | Olf. |  |  |  |  |  |  |
| Motor |  | 0.0048 | 0.85 |  |  |  |  |  |  |
| PFC | 0.0048 |  | 0.0016 |  |  |  |  |  |  |
| Olfactory | 0.85 | 0.0016 |  |  |  |  |  |  |  |

**Supplemental Table 4**. Bootstrap (top) and generalized linear mixed-effects model (bottom), *p*-values from pairwise comparisons with Bonferroni correction (related to Fig. 5).
