## Supplemental Table 5 for "A stable, distributed code for cue value in mouse cortex during reward learning"

| **Value cells (prop cue), Bonferroni corrected p-values for pairwise region comparisons (Fig. S10A-D, left)** | | | | | | | | | |
| --- | --- | --- | --- | --- | --- | --- | --- | --- | --- |
|  | ALM | ACA | FRP | PL | ILA | ORB | DP | TTd | AON |
| ALM |  | 1 | 1 | 1 | 1 | 1 | 0.17 | 0.0187 | 0.0549 |
| ACA | 1 |  | 1 | 1 | 0.17 | 0.20 | 0.0094 | 0.0010 | 0.0043 |
| FRP | 1 | 1 |  | 1 | 1 | 1 | 1 | 0.22 | 0.35 |
| PL | 1 | 1 | 1 |  | 0.79 | 0.45 | 0.0229 | 0.0040 | 0.0229 |
| ILA | 1 | 0.17 | 1 | 0.79 |  | 1 | 1 | 0.66 | 1 |
| ORB | 1 | 1 | 1 | 0.45 | 1 |  | 1 | 0.16 | 0.34 |
| DP | 0.17 | 0.0094 | 1 | 0.0229 | 1 | 1 |  | 1 | 1 |
| TTd | 0.0187 | 0.0010 | 0.22 | 0.0040 | 0.66 | 0.16 | 1 |  | 1 |
| AON | 0.0549 | 0.0043 | 0.35 | 0.0229 | 1 | 0.34 | 1 | 1 |  |
|  | Motor | PFC | Olfactory |  |  |  |  |  |  |
| Motor |  | 0.78 | 6.80E-07 |  |  |  |  |  |  |
| PFC | 0.78 |  | 6.27E-09 |  |  |  |  |  |  |
| Olfactory | 6.80E-07 | 6.27E-09 |  |  |  |  |  |  |  |
| **Value-like cells (prop cue), Bonferroni corrected p-values for pairwise region comparisons (Fig. S10A-D, right)** | | | | | | | | | |
|  | ALM | ACA | FRP | PL | ILA | ORB | DP | TTd | AON |
| ALM |  | 1 | 1 | 1 | 1 | 1 | 0.57 | 1 | 0.42 |
| ACA | 1 |  | 1 | 1 | 1 | 1 | 1 | 1 | 1 |
| FRP | 1 | 1 |  | 1 | 1 | 1 | 1 | 1 | 1 |
| PL | 1 | 1 | 1 |  | 1 | 1 | 1 | 1 | 1 |
| ILA | 1 | 1 | 1 | 1 |  | 1 | 1 | 1 | 1 |
| ORB | 1 | 1 | 1 | 1 | 1 |  | 1 | 1 | 1 |
| DP | 0.57 | 1 | 1 | 1 | 1 | 1 |  | 1 | 1 |
| TTd | 1 | 1 | 1 | 1 | 1 | 1 | 1 |  | 1 |
| AON | 0.42 | 1 | 1 | 1 | 1 | 1 | 1 | 1 |  |
|  | Motor | PFC | Olfactory |  |  |  |  |  |  |
| Motor |  | 0.37 | 0.092 |  |  |  |  |  |  |
| PFC | 0.37 |  | 0.63 |  |  |  |  |  |  |
| Olfactory | 0.092 | 0.63 |  |  |  |  |  |  |  |

**Supplemental Table 5**. Generalized linear mixed-effects model, *p*-values from region contrasts (related to Fig. S10).
