## Supplemental Table 1 for "A stable, distributed code for cue value in mouse cortex during reward learning"

| **Cue cells, Bonferroni corrected p-values for pairwise region comparisons (Fig. 2G, left)** | | | | | | | | | |
| --- | --- | --- | --- | --- | --- | --- | --- | --- | --- |
|  | ALM | ACA | FRP | PL | ILA | ORB | DP | TTd | AON |
| ALM |  | 0.1580 | 1 | 1.68E-07 | 4.17E-10 | 6.19E-25 | 3.04E-17 | 7.06E-18 | 8.38E-17 |
| ACA | 0.1580 |  | 1 | 1 | 1 | 0.0654 | 0.0457 | 0.0121 | 0.0191 |
| FRP | 1 | 1 |  | 0.4334 | 0.0234 | 9.27E-09 | 3.05E-06 | 7.44E-07 | 1.13E-07 |
| PL | 1.68E-07 | 1 | 0.4334 |  | 1 | 2.86E-05 | 0.0002 | 3.28E-05 | 0.0004 |
| ILA | 4.17E-10 | 1 | 0.0234 | 1 |  | 0.2634 | 0.1894 | 0.0446 | 0.1065 |
| ORB | 6.19E-25 | 0.0654 | 9.27E-09 | 2.86E-05 | 0.2634 |  | 1 | 1 | 1 |
| DP | 3.04E-17 | 0.0457 | 3.05E-06 | 0.0002 | 0.1894 | 1 |  | 1 | 1 |
| TTd | 7.06E-18 | 0.0121 | 7.44E-07 | 3.28E-05 | 0.0446 | 1 | 1 |  | 1 |
| AON | 8.38E-17 | 0.0191 | 1.13E-07 | 0.0004 | 0.1065 | 1 | 1 | 1 |  |
| **Lick cells, Bonferroni corrected p-values for pairwise region comparisons (Fig. 2G, right)** | | | | | | | | | |
|  | ALM | ACA | FRP | PL | ILA | ORB | DP | TTd | AON |
| ALM |  | 1 | 0.1505 | 0.0011 | 3.46E-06 | 2.51E-06 | 8.87E-05 | 0.0018 | 0.0434 |
| ACA | 1 |  | 1 | 1 | 0.4797 | 1 | 0.1987 | 1 | 1 |
| FRP | 0.1505 | 1 |  | 1 | 1 | 1 | 1 | 1 | 1 |
| PL | 0.0011 | 1 | 1 |  | 1 | 1 | 0.6895 | 1 | 1 |
| ILA | 3.46E-06 | 0.4797 | 1 | 1 |  | 1 | 1 | 1 | 1 |
| ORB | 2.51E-06 | 1 | 1 | 1 | 1 |  | 1 | 1 | 1 |
| DP | 8.87E-05 | 0.1987 | 1 | 0.6895 | 1 | 1 |  | 1 | 1 |
| TTd | 0.0018 | 1 | 1 | 1 | 1 | 1 | 1 |  | 1 |
| AON | 0.0434 | 1 | 1 | 1 | 1 | 1 | 1 | 1 |  |
| **Both cells, Bonferroni corrected p-values for pairwise region comparisons (Fig. S6B, right)** | | | | | | | | | |
|  | ALM | ACA | FRP | PL | ILA | ORB | DP | TTd | AON |
| ALM |  | 0.0637 | 1 | 0.0005 | 1 | 0.1383 | 0.1388 | 2.29E-06 | 7.50E-05 |
| ACA | 0.0637 |  | 1 | 1 | 1 | 1 | 1 | 1 | 1 |
| FRP | 0.2811 | 1 |  | 1 | 1 | 1 | 1 | 1 | 1 |
| PL | 0.0005 | 1 | 1 |  | 1 | 1 | 1 | 1 | 1 |
| ILA | 1 | 1 | 1 | 1 |  | 1 | 1 | 0.0250 | 0.20 |
| ORB | 0.1383 | 1 | 1 | 1 | 1 |  | 1 | 0.0398 | 0.0574 |
| DP | 0.1388 | 1 | 1 | 1 | 1 | 1 |  | 1 | 1 |
| TTd | 2.29E-06 | 1 | 1 | 1 | 0.0250 | 0.0398 | 1 |  | 1 |
| AON | 7.50E-05 | 1 | 1 | 1 | 0.20 | 0.0574 | 1 | 1 |  |

**Supplemental Table 1**. Generalized linear mixed-effects model, *p*-values from region contrasts (related to Fig. 2).
